# Multi-omics Comparison among Populations of Three Plant Sources of Amomi Fructus

**DOI:** 10.1101/2023.01.27.525993

**Authors:** Xinlian Chen, Shichao Sun, Xiaoxu Han, Cheng Li, Bao Nie, Zhuangwei Hou, Jiaojiao Ji, Xiaoyu Han, Lixia Zhang, Jianjun Yue, Depo Yang, Li Wang

## Abstract

Amomi Fructus (Sharen, AF) is a traditional Chinese medicine (TCM) from three source species (or subspecies) including *Wurfbainia villosa* var. *villosa* (WVV), *W. villosa* var. *xanthioides* (WVX) or *W. longiligularis* (WL). Among them, WVV has been transplanted from its top-geoherb region Guangdong to its current main production area Yunnan for more than 50 years in China. However, the genetic and transcriptomic differentiation among multiple AF source (sub)species and between the origin and transplanted populations of WVV is unknown. In our study, the observed overall higher expression of terpenoid biosynthesis genes in WVV than that of WVX supplied possible evidence for the better pharmacological effect of WVV. We also screened ten candidate *borneol dehydrogenase* (*BDH*) genes that potentially catalyzed borneol into camphor in WVV. The *BDH* genes may experience independent evolution after acquiring the ancestral copies and the followed tandem duplications might account for the abundant camphor content in WVV. Furthermore, four populations of WVV, WVX and WL are genetically differentiated and the gene flow from WVX to WVV in Yunnan contributed to the increased genetic diversity in the introduced population (WVV-JH) compared to its top-geoherb region (WVV-YC), which showed the lowest genetic diversity and might undergo genetic degradation. In addition, *TPS* and *BDH* genes were selected among populations of multiple AF source (sub)species and between the top-geoherb and non-top-geoherb regions, which might explain the metabolite difference of these populations. Our findings provide important guidance for the conservation, genetic improvement, industrial development of the three source (sub)species, and identifying top-geoherbalism with molecular markers and proper clinical application of AF.

## INTRODUCTION

Top-geoherbalism, similar to “*Daodi*” in China and “Provenance” or “Terroir” in Europe, refers to traditional herbs grown in certain native ranges with better quality and efficacy than those grown elsewhere^1,2^, which reflects the difference of the component and abundance of secondary metabolites among populations, resulting from multiple factors, such as genetic elements, environmental factors, cultural processing etc^3-5^. The varied efficacy of traditional medicine with multiple sources results in the mixture of the inferior and the superior in the market, which inhibits the standardization and internalization of traditional medicine.

One special case of top-geoherbalism comes from the legally accredited multiple source plant species (or subspecies) for a single medicine. The abundance of chemical components, often listed as the quality control of traditional Chinese medicine (TCM), in different origin species are regularly different, and thus the pharmacological effect of each origin varies^6-9^, although they are used as the same TCM clinically. For example, Ephedrae herba is the dry herbaceous stem of *Ephedra sinica, E. intermedia* or *E. equisetina*^10^. But total alkaloid abundance in *E. equisetina* with the greatest acute toxicity is higher than that in the first two species^11,12^. The main alkaloid accumulated in *E. sinica* with the best effect of relaxation and cough relieving was Ephedrine, while in *E. intermedia* was (+)-pseudoephedrine^11,13,14^. Another common case of top-geoherbalism lies in that different populations of the same species demonstrate distinguished pharmacological efficacy. For example, Tan et al. identified chemical markers β-ocimene, α-pinene, 3-methylbutanal, heptanes, butanal for distinguishing Radix *Angelica sinensis* from its top-geoherb regions with superior clinical practice compared with that from non-top-geoherb regions^15^.

Amomi Fructus (Sharen, AF) provides an ideal system to investigate the top-geoherbalism, as it is legally recorded from multiple (sub)species and the varied populations of the same species exhibit distinct biochemical components and abundance. In Chinese Pharmacopoeia, AF is described as the dry and mature fruit of *Wurfbainia villosa* var. *villosa* (Lour.) Škorničk. & A.D.Poulsen (WVV, Figure 1A), *Wurfbainia villosa* var. *xanthioides* (Wall. ex Kuntze) Škorničk. & A.D.Poulsen (WVX, Figure 1A) or *Wurfbainia longiligularis* (T.L.Wu) Škorničk. & A.D.Poulsen (WL), in the Zingiberaceae family^10^. It is one of the four most important Southern China Medicines, and a dual-purpose commodity for medicine and food^16-18^. It is well-known for its efficacy of soothing the fetus, stopping diarrhea and appetizing^10^, and the effective components are volatile oil, mainly terpenes, including monoterpenes (bornyl acetate, borneol, camphor, myrcene, limonene, α-terpinene, etc.) and sesquiterpenes (germacrene, bicyclogermacrene, α-copaene, α-santalol, etc.)^19^.

**FIGURE 1.**
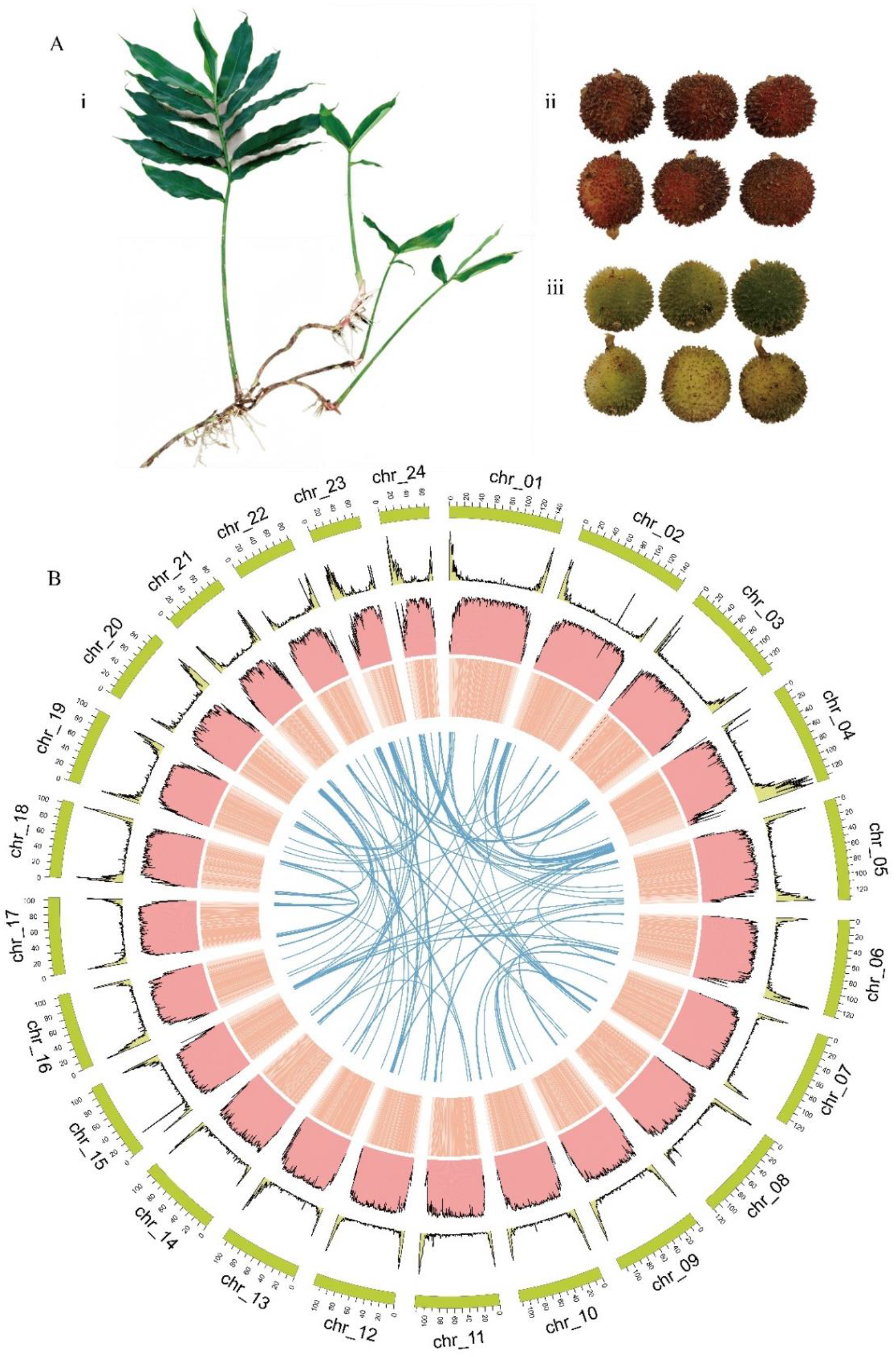
Overview of WVV genome assembly and genomic features. (A) Morphological characteristics of WVV and WVX. (i) Plant and (ii) fruits of WVV and (iii) fruits of WVX. (B) Distribution of WVV genomic features. The circos represented synteny, GC content, TE distribution, gene density and karyotypes from inside to outside, respectively. All these genomic features were calculated with 500 kb non-overlapped sliding windows.

Profound difference has been observed in the abundance of volatile metabolites of the three plant (sub)species of AF^20-22^, which is reflected in the different standards for the abundance of volatile oil of the three source (sub)species in Chinese Pharmacopoeia. The volatile oil abundance in the seed of WVV and WVX are required to be no less than 3.0% (ml/g), and no less than 1.0% (ml/g) in WL^10^. The main volatile component of WVV and WL is bornyl acetate, and camphor for WVX^20-22^. From the clustering analysis of volatile metabolites identified via High Performance Liquid Chromatography (HPLC), WVV and WL showed similar patterns, and they were clearly differentiated from WVX^23^. AF in the market is sometimes a mixture of WVV and WVX, whose fruit color is the same as that of WVV after processing. Thus, the genetic characteristics of the three (sub)species, including the genomic variation and the gene expression underlying the difference of volatile compounds, awaits further clarification for the molecular identification of medicinal sources.

In addition, the top-geoherb region of WVV is Yangchun, Guangdong province, China^18^. However, WVV in Yangchun must be artificially pollinated, and thus demonstrates poor yields. Since 1960s, WVV has been gradually introduced to Guangxi, Fujian and Yunnan province, where the natural pollination happens via local insects and it greatly increases its yield. Currently, the production of WVV from Yunnan accounts for more than 80% of the market share, but the price was 20 times lower than that from Yangchun owing to the label of “top-geoherbalism” products^18^. However, it is controversial with regard to the efficacy of WVV from Guangdong and Yunnan provinces. Some researchers found there was no significant difference in the pharmacological activities of WVV from the two localities^24^. While some studies found that the comprehensive quality score such as abundance of volatile oil and bornyl acetate of WVV in Guangdong was the highest among WVV from other places^25,26^. Given that the genetic separation of WVV from Guangdong and Yunnan is only over 50 years, it is obscure concerning the genetic variation of WVV from the top-geoherb and main production areas.

Genetic studies have been advanced by a recent release of the genome of WVV^27^, and a large number of *terpene synthesis* (*TPS*) genes have been screened and verified, revealing parts of biosynthetic pathway of terpenes in WVV^27-32^. The identified genes included linalool synthase gene (*AvTPS2*), α-santalene and α-bergimonene synthase gene (*AvTPS15*)^33^, α-pinene and β-pinene synthase gene (*AvPS*)^34^. The monoterpene bornyl acetate is one of the most significant characteristic substances in WVV. Previous study unveiled that *WvBAT3* and *WvBAT4* might be the two key *borneol acetyltransferases* (*BAHD*) for its synthesis in the seeds of WVV^27^. However, the genes catalyzing borneol to camphor have not been resolved in WVV. According to the chemical structures of borneol and camphor, this oxidation step is possibly catalyzed by the *borneol dehydrogenase* (*BDH*). *BDH* genes have been cloned and functionally verified in several species, such as *CcBDH3* in *Cinnamomum camphora*^35^, *LiBDH* in *Lavandula x intermedia*^36^, *AaBHD* in *Artermisia annua*^37^. They belong to *short-chain dehydrogenases/reductases* (*SDR*s) subfamily^36,38^, constituting a large family of NAD(P)(H)-dependent oxidoreductases, sharing sequence motifs and displaying similar mechanisms^39,40^. Taken together, *BDH* genes catalyzing borneol to camphor in WVV await further exploration.

Here we *de novo* assembled a chromosome-level genome of WVV from its top-geoherb location and conducted comprehensive comparison among populations of the three plant (sub)species genetically and transcriptionally and between the top-geoherb and main production areas of WVV, and we aimed to address: 1) the expressional difference of terpenes related genes between WVV and WVX; 2) the candidate *BDH* genes catalyzing borneol into camphor in WVV; 3) genetic differentiation among populations of WVV, WVX and WL. Our findings will provide important guidance for the conservation, genetic improvement of AF source (sub)species, and identifying top-geoherbalism and proper clinical application of AF.

## RESULTS

### Genome Sequencing, Assembly and Annotation of *W. villosa* var. *villosa*

The genome size of WVV was estimated as ∼2.62 Gb with flow cytometry, which gave us a rough hint of the amount of sequencing data to produce a good quality *de novo* assembled genome. In total, we obtained 157.07 Gb (∼55.50X) High-fidelity (Hi-fi) long reads and 286.28 Gb (∼101.16X) high-throughput chromosome conformation capture (Hi-C) short reads. This allowed us to obtain a haplotype-resolved chromosome-level genome assembly. The assembly resulted in haplotype 1 of 2,901 contigs (N50 = 8.30 Mb), with a total size of 2.83 Gb (Table 1). The genome size of haplotype 2 was 2.77 Gb with contig N50 of 7.01 Mb. Subsequently, the larger haplotype genome, haplotype 1, was chosen for the following analysis, and its contig sets were anchored based on Hi-C contacts. It was anchored to 24 pseudochromosomes with the scaffolding rate of 94.2% (Figure S1). The final assembled WVV genome was 2.83 Gb (Table 1). The scaffold N50 was 112.82 Mb. The length of 24 pseudochromosomes was from 151,371,763 (chr_01) to 66,131,911 bp (chr_24) (Figure 1B and Table S1). The genome size of WVV was almost the same as the published WVV genome size (2.80 Gb)^17^ and 1.4∼2.9 times larger than that of other Zingiberaceae species^41-44^. The average GC content of WVV genome was 40.73% (Table 1). To test the quality of the WVV genome assembly, RNA-seq paired-end reads were mapped to the assembled genome, with mapping rates of 92.08-96.61%. In addition, Benchmarking Universal Single-Copy Orthologs assessment (BUSCO) analysis showed that the assembled genome covered 99.8% of the viridiplantae orthologous gene set (Table 1). Taken together, the above evidence suggested the completeness of the assembled genome.

**TABLE 1.** Assembly and annotation statistics of the genome.

| Parameter | WVV |
| --- | --- |
| **Contig** |  |
| Total assembly size (bp) | 2,834,193,068 |
| Total contig number | 2,901 |
| Maximum contig length (bp) | 56,657,728 |
| Contig N50 length (bp) | 8,295,749 |
| Contig N90 length (bp) | 1,378,773 |
| **Scaffold** |  |
| Total assembly size (bp) | 2,834,739,068 |
| Total scaffold number | 2,605 |
| Maximum scaffold length (bp) | 151,371,763 |
| Scaffold N50 (bp) | 112,824,243 |
| Scaffold N90 (bp) | 68,159,171 |
| GC content (%) | 40.73 |
| Complete BUSCOs (%) | 99.8 |
| **Annotation** |  |
| Gene number | 50,473 |
| Repeat content (%) | 85.67 |
| GO scale (%) | 34.08 |
| KEGG scale (%) | 34.28 |

Combining *ab initio* prediction, orthologous protein and transcriptomic data, we annotated 50,473 coding genes, of which 39,556 genes were located on 24 pseudochromosomes (Table 1 and Table S1). The average gene density was one gene per 56.15 kb, with the genes unevenly distributed, being more abundant towards the chromosomal ends (Figure 1b). The average length of coding sequences of the predicted genes was 907.49 bp, with an average of 3.79 exons per gene (Table S2). Approximately 77.98% of the protein-coding genes were functionally annotated by searching SwissProt, Kyoto Encyclopedia of Genes and Genomes (KEGG), Pfam and Gene Ontology (GO) databases (Table S3). Transposable element (TE) sequences comprised 85.67% (2.43 G) of WVV genome. Among them, LTR (Long-terminal repeat) was the most abundant repetitive type (79.68%) and *Copia* was the largest group in the LTR, accounting for 50% of the entire genome (Table S4).

### Comparison of Expression Levels of Terpenoid Biosynthetic Genes between *W. villosa* var. *villosa* and *W. villosa* var. *xanthioides*

The abundance of certain secondary metabolites between WVV and WVX were obviously different according to previous studies (Figure 2A)^21^. The abundance of terpinolene, α-pinene, camphene and bornyl acetate was higher in WVV, while linalool, β-myrcene, borneol and camphor were richer in WVX. The terpenoid biosynthesis in WVV and WVX mainly included three types of genes: *TPS, BAHD* and *BDH*. To investigate the pattern of gene expression affecting the abundance of volatile compounds between the two subspecies, we analyzed the expression of genes involved in multiple terpenes biosynthesis in various tissues and fruit developmental stages between WVV and WVX (see Materials and Methods; Figure 2).

**FIGURE 2.**
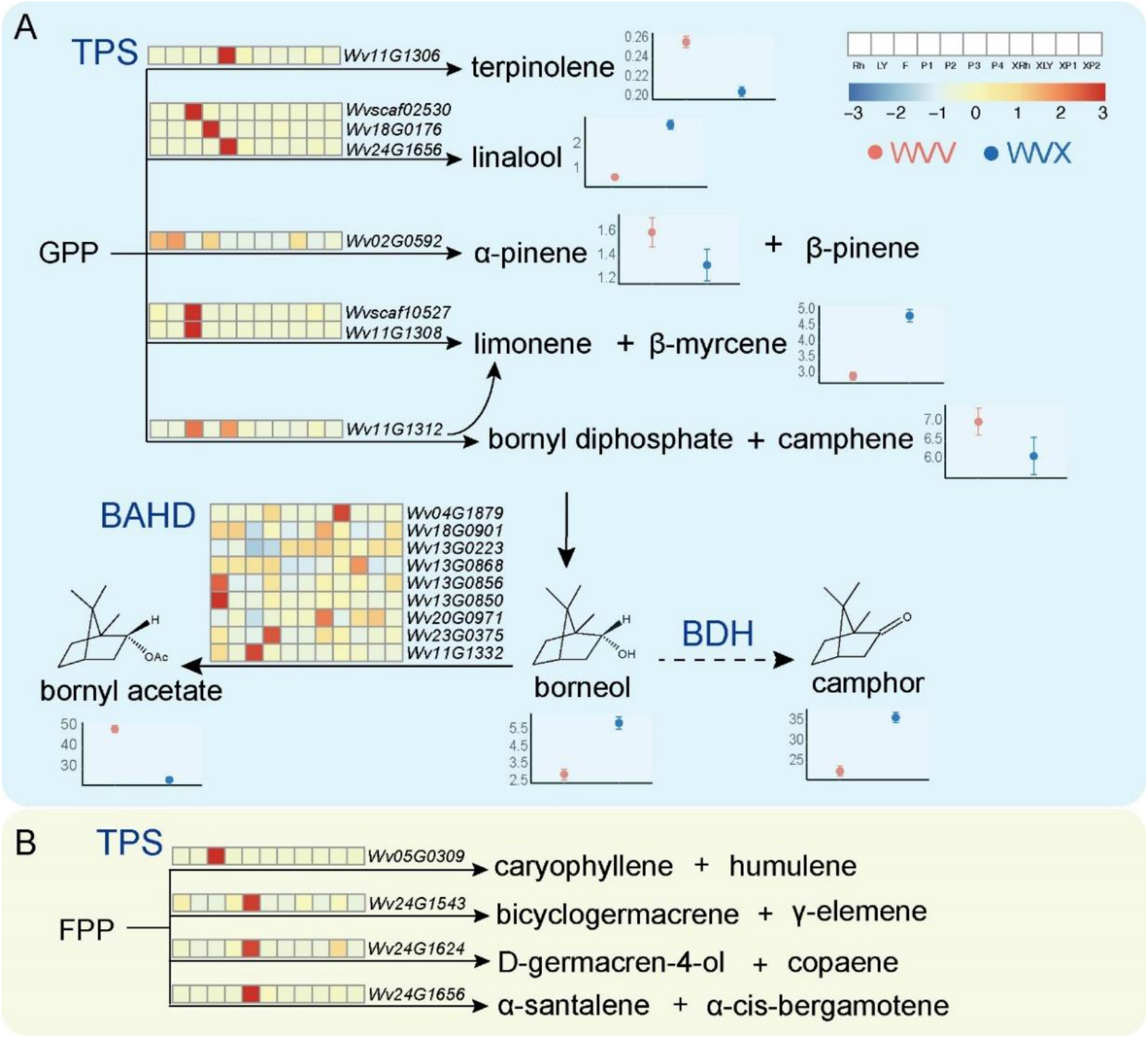
The heatmap of terpene synthesis-related gene expressions from different tissues and fruit developmental stages in WVV and WVX. The heatmap represented rhizomes (Rh), young leaves (LY), flowers (F), 10-day after flowering (P1), 30-day after flowering (P2), 60-day after flowering (P3), 90-day after flowering (P4) of WVV, and rhizomes (XRh), young leaves (XLY), 75-day after flowering (XP1), 90-day after flowering (XP2) of WVX from left to right. On the right or below of the chemical structure was the difference in the relative content of each volatile oil in the fruits reported in Ao et al.^21^ The x-axis showed WVV (red) and WVX (blue). The y-axis represented the relative content. (A) Monoterpene synthesis. (B) Sesquiterpene synthesis.

Subsequently, we constructed the maximum-likelihood (ML) tree with *TPS* genes from WVV genome and the functionally verified *TPS* genes in previous study^17^. Based on the ML tree, 11 *TPS* genes, clustered with the verified ones, were selected (Figure S2). The transcriptome data were generated for the rhizomes (Rh), young leaves (LY), flowers (F), fruits of 10-day after flowering (P1), 30-day after flowering (P2), 60-day after flowering (P3), and 90-day after flowering (P4) of WVV and rhizomes (XRh), young leaves (XLY), fruits of 75-day after flowering (XP1), and 90-day after flowering (XP2) of WVX to compare the expressional difference of the two subspecies. Overall, the expression level of these *TPS* genes in WVV was relatively higher than that of WVX (Figure 2) and high expression of *TPS* genes did not always coincide with metabolite abundance trends in WVV and WVX. For the monoterpene biosynthesis genes, the highest expression level of *Wv11G1306* was observed at P2 stage, consistent with the higher abundance of terpinolene in the fruits of WVV (Figure 2A). High expressions of *TPS* genes *Wv18G0176* and *Wv24G1656* (for linalool) appeared in P1 and P2 stages, separately, contrast with the trend of the abundance of linalool in WVV and WVX. *Wv02G0592, Wvscaf10527, Wv11G1308* and *Wv11G1312* were all bi- or multi-functional enzyme genes based on verified homologous genes^17^. We only knew the abundance trends of α-pinene, β-myrcene, and camphene of WVV and WVX, not that of β-pinene, limonene and bornyl diphosphate^21^. While gene expression level of *Wv02G0592* (for α-pinene and β-pinene), *Wv11G1312* (for limonene, bornyl diphosphate, camphene), *Wvscaf10527* and *Wv11G1308* (for limonene and β-myrcene) were low expressed in fruits of both subspecies. In addition, for sesquiterpenoids, *Wv24G1543* catalyzing the substrate into bicyclogermacrene and γ-elemene, *Wv24G1624* into D-germacren-4-ol and copaene, and *Wv24G1656* into α-santalene and α-cis-bergamotene were all expressed at a high level at P2 stage (Figure 2B). The inconsistency between expression level of *TPS* genes and metabolite abundance trends could be related with multiple factors: 1) the absent collection of younger fruit in WVX, missing the high expression period; 2) the different source of expression and secondary metabolite data; 3) the potential transportation of secondary metabolites from its biosynthetic organs to other tissues.

As for the three most significant chemical substances in WVV and WVX, with the borneol as the same substrate, the bornyl acetate was produced by *BAHD* genes while the camphor by *BDH* genes. We located nine *BAHD* genes in this genome according to experimentally verified *BAHD*s^27^ (Figure 2A). The higher expression levels of *Wv18G0901, Wv13G0223* and *Wv20G0971* were at P4 stage, and *Wv23G0375* at P1 stage, may be the key candidate genes (Figure 2A).

To further explore the regulation of biosynthetic genes, 2,781 TFs were identified in WVV genome. To discern TFs regulating bornyl acetate biosynthesis in WVV, we constructed a gene co-expression network. We further analyzed TFs regulating the above-mentioned *BAHD* genes. 189, 278, 81, 203, 92, 1, and 276 TFs were involved in regulating *Wv04G1879, Wv11G1332, Wv13G0223, Wv13G0850, Wv13G0856, Wv13G0868*, and *Wv23G0375* genes, respectively (Figure S3 and Table S7). The well-known MYB and WRKY TFs were found to be involved in regulating bornyl acetate biosynthesis (Figure S3). Previous studies showed that MYB and WRKY TFs participated in regulating the synthesis of terpenes^45-48^, and in particular, WRKY was found to play important roles in bornyl acetate biosynthesis^31^. This analysis provided a list of TF candidates for further functional verification in bornyl acetate biosynthesis.

### Candidate *BDH* Genes and their Potential TFs Catalyzing Borneol into Camphor

To investigate the evolutionary relationships among *BDH* genes and identify candidate genes in WVV, we constructed a phylogenetic tree. The tree contained BDH proteins from one monocot (WVV), one gymnosperm (*Taxus chinensis*, as the outgroup), one magnoliids (*C. camphora*), and three eudicots (*Salvia officinalis, A. annua, Rosmarinus officinalis*), and the numbers of BDHs in the above-mentioned species were 29, 14, 16, 19, 20, and 18, respectively. Nine experimentally verified BDH proteins from five species were also included (Figure 3A; Supplementary data 1; Table S5). In total, the tree included 125 BDHs from seven species, among which there was only one from *L. intermedia*, LiBDH, owing to the absence of its genome.

**FIGURE 3.**
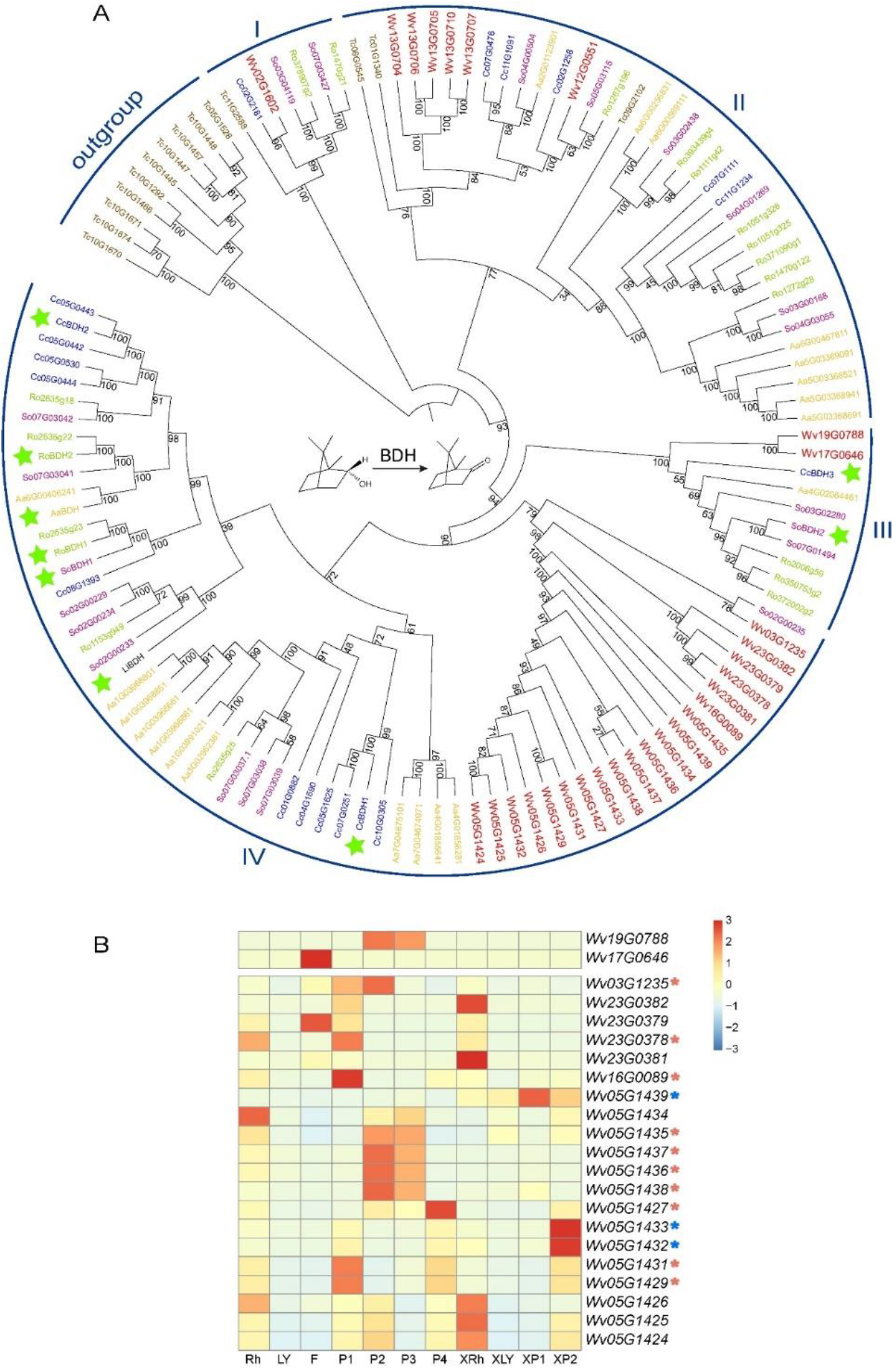
Phylogenetic tree of BDH homologous proteins in various species. (A) The phylogenetic tree of BDH genes. Wv: WVV. Cc: *Cinnamomum camphora*. So: *Salvia officinalis*. Tc: *Taxus chinensis*. Aa: *Artemisia annua*. Ro: *Rosmarinus officinalis*. The green stars marked the verified BDH enzymes. The numbers on the branches showed the bootstraps. (B) The heatmap of expression level of candidate *BDH* genes identified in clades III and IV in different tissues and developmental periods of WVV. Red asterisks indicated ten *BDH* genes, which were highly expressed in WVV fruit developmental stages, and blue asterisks showed three *BDH* genes, which were highly expressed in WVX fruit developmental stages.

Except the 10 BDHs of the outgroup, we divided the remaining members into four clades from I to IV and the bootstrap of each clade was over 75 (Figure 3A). The BDH protein numbers from clades I to IV were 7, 36, 10, and 62, respectively. Only clade I and clade II had BDHs from gymnosperm, suggesting that clades III and IV may have originated from these two clades and BDHs in clades I and II appeared before the divergence between gymnosperm and angiosperm. BDHs of each species were dispersed in different clades with clades I to IV containing 5, 6, 5, and 6 BDHs, respectively, suggesting that these species underwent independent evolution after acquiring ancestral copies. Clades I to IV included 1, 6, 2, and 20 BDHs of WVV, respectively (Figure 3A). Clade III comprised Wv19G0788, Wv17G0646 and two validated enzymes CcBDH3 and SoBDH2. Clade IV consisted of 20 BDHs in WVV and seven experimentally validated BDHs, CcBDH1, LiBDH, SoBDH1, RoBDH1, AaBDH, RoBDH2, and CcBDH2. Thus, we speculated the 22 BDHs in clades III and IV were more likely candidate enzymes that catalyzed the dehydrogenation of borneol to form camphor in WVV. Interestingly, we found the WVV BDHs of clade IV were compactly distributed on chromosomes 23 and 5, suggesting couple of tandem duplication events, which were also reflected in the BDHs from *T. chinensis, A. annua*, and *C. camphora* (Figure 3A).

To further narrow down the candidate *BDH*s in WVV, we examined expression levels of the 22 candidate *BDH*s in various tissues and fruit developmental stages (Figure 3, Table S6). For clade III, *Wv19G0788* was highly expressed in both P2 and P3 and *Wv17G0646* was lowly expressed in any fruit stage. However, the expression level of *Wv19G0788* and *Wv17G0646* in all tissues was almost 0, thus they were not likely candidate genes (Table S6). For clade IV (20 WVV *BDH*s), ten genes (*Wv03G1235, Wv23G0378, Wv16G0089, Wv05G1435, 1436, 1437, 1438, 1427, 1431, 1429*) were at least highly expressed in one fruit stage, more likely candidate *BDH* genes in the fruits of WVV (Table S6). In addition, *Wv05G1439, Wv05G1433*, and *Wv05G1432* were only highly expressed in the fruit stages of WVX, but not in WVV. The expressional differences of the two groups of genes may be the underlying reason for the difference in camphor abundance between WVV and WVX.

We further explored the expressional regulation of the candidate genes. *BDH* genes, *Wv03G1235, Wv05G1427, 1429, 1431, 1435, 1436, 1437, 1438, Wv16G0089* and *Wv23G0378*, were regulated by 80, 18, 159, 159, 11, 102, 102, 115, 292 and 292 TFs, respectively (Figure S4 and Table S8), including bHLH, WRKY, NAC, and MYB families, which were reported to be crucial in plant growth and development, stress resistance, and secondary metabolism^49,50^. Remarkably, tandem duplicated *BDH* genes (*Wv05G1429, 1431, 1435, 1436, 1437* and *1438*) were regulated by the same TFs, suggesting the co-regulated expression pattern of tandem duplicated genes. For example, 159 TFs synchronously regulated *Wv05G1429* and *1431*. While *Wv05G1435, 1436, 1437* and *1438* were simultaneously regulated by GRAS (*Wv02G0982*), ZF-HD (*Wv04G3465*), bZIP (*Wv09G0998*), NF-YB (*Wv13G0841*), ERF (*Wv20G1381*), MYB (*Wv02G0066, Wv08G0553* and *Wv08G0729*), etc. Those TFs and the tandemly duplicated candidate *BDH* genes especially await further experimental verification.

### Re-sequencing of Amomi Fructus populations

To explore genetic variation of the three source (sub)species of AF, leaf tissues from 39 accessions, which contained 20 samples of WVV (10 WVV-YC from Yangchun, Guangdong and 10 WVV-JH from Jinghong, Yunnan), 10 WVX and nine WL, were used for the construction of 150-bp paired-end libraries, and then sequenced, separately. In total, 1.94 Tb (10.0X-29.5X) raw data were obtained, and the average sequencing depth was 17.5X.

The re-sequenced individuals were collected from three geographic locations (Figure 4A). After mapping to the assembled WVV genome, we identified 24,854,114 putative single-nucleotide polymorphisms (SNPs). We characterized the genetic relationships among WVV-YC, WVV-JH, WVX and WL with neighbor-joining (NJ) phylogenetic tree (Figure 4C), as well as principal component analysis (PCA) (Figure S5). The three origin (sub)species were mainly divided into four genetic groups, corresponding to the four sampled populations. Based on the filtered SNP dataset (see “Methods”), the optimal number of populations in the STRUCTURE analysis was *K* = 3 (Figure S6). At *K* = 3, one cluster was consisted of all the WL samples, the second included seven WVX, and the third contained 17 WVV (ten from YC, seven from JH), with three individuals of WVV-JH and three of WVX exhibiting admixture (the percentage of the minor component was bigger than 50%; Figure 4E). At *K* = 4, WVV was further divided into a WVV-YC and a WVV-JH group, the former comprising ten individuals from Yangchun (the top-geoherb location of WVV), the latter comprising seven individuals from Jinghong, which were introduced from Yangchun at around 50 years ago.

**FIGURE 4.**
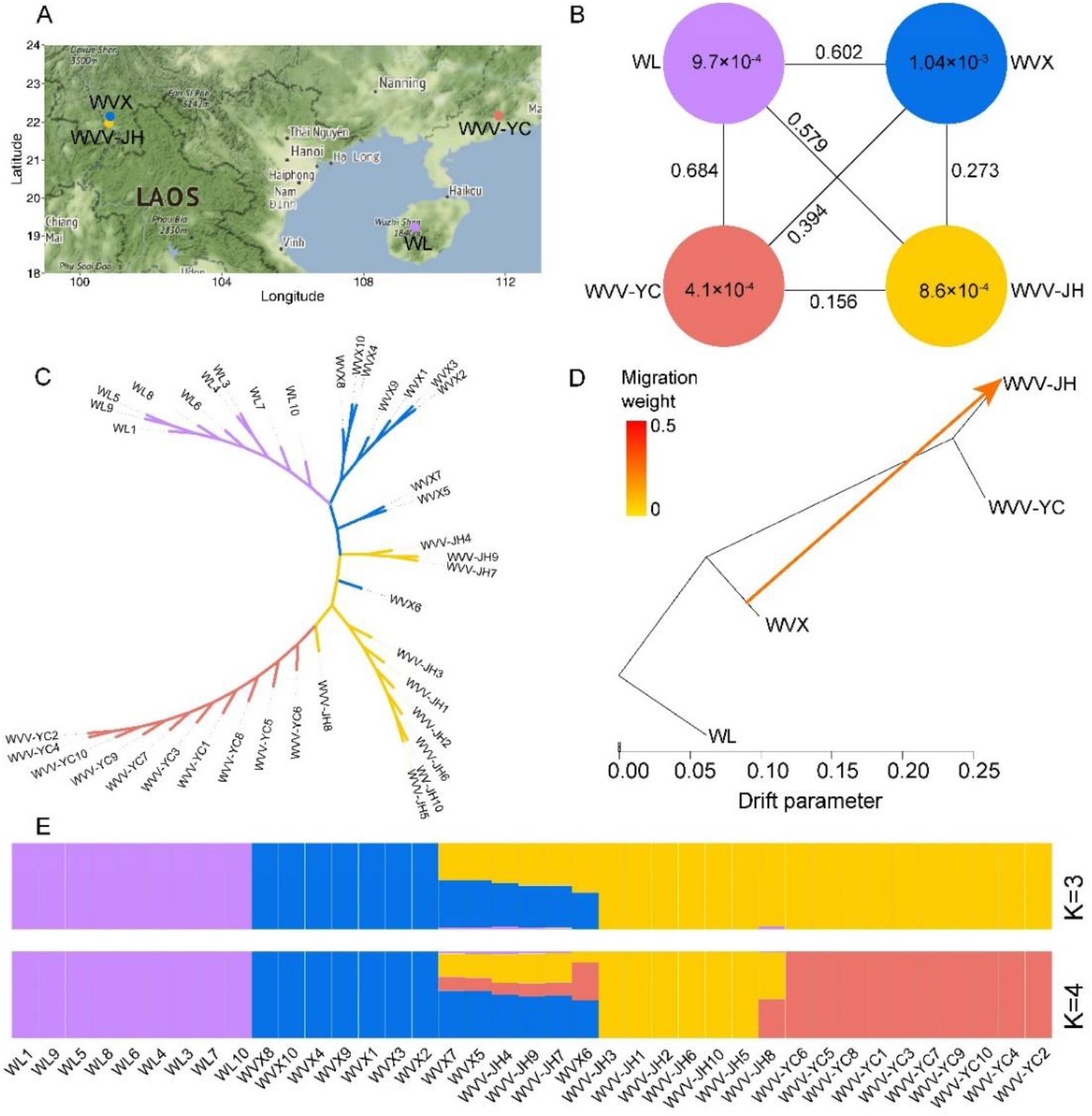
Population genetic analysis of three source (sub)species of Amomi Fructus. The color code for the populations is consistent in the figure: purple (WL), red (WVV-YC), blue (WVX) and orange (WVV-JH). (A) Geographical distribution of four populations, including WVV-YC, WVV-JH, WVX, and WL. (B) Population differentiation *F*ST among populations and the nucleotide diversity π of each population based on 24,854,114 SNPs. (C) A NJ tree of 39 accessions based on 160,967 high-quality SNPs. (D) The gene flow from WVX to WVV-JH identified in the TreeMix analyses. The arrow indicates the migration direction. (E) Population structure analyses showed the differentiation of 39 accessions.

To study the genetic differentiation among and within the three (sub)species of AF, we estimated population differentiation *F*_ST_ between populations and the nucleotide diversity π within species. Based on the high-density SNP data, π of the WVX population was estimated to be 1.04×10^−3^, which is slightly higher than that of the WL (π = 9.7×10^−4^) and WVV-JH (π = 8.6×10^−4^) populations, and these three populations exhibited considerably higher genetic diversity than that of the WVV-YC (π = 4.1×10^− 4^) (Figure 4B), suggesting the genetic bottleneck of the narrowly distributed top-geoherbalism population. It was worth noting that the nucleotide diversity in the WVV-JH (π = 8.6×10^−4^) was more than twice of that in the origin location of WVV-YC (π = 4.1×10^−4^), which could result from the gene flow with the locally occurring WVX (Figure 4D). To test this hypothesis, we removed the three admixed individuals of WVV-JH, the diversity of WVV-JH was reduced to 5.0×10^−4^, but was still higher than that in WVV-YC. It suggested that human-mediated pollination could not fully compensate the occurrence of inbreeding in WVV-YC owing to the small population size, and the introduced population in Yunnan, independent of human-mediated pollination, might have a higher rate of outcrossing and thus restore the genetic diversity. The highest *F*_ST_ (0.684) was between WL and WVV-YC, followed by that between WL and WVX (*F*_ST_ = 0.602), between WL and WVV-JH (*F*_ST_ = 0.579), as WL and WVV/WVX were obviously different species. The middle rank was between WVX and WVV-YC (*F*_ST_ = 0.394), WVX and WVV-JH (*F*_ST_ = 0.273). As WVV-JH and WVX were from similar geographical environment, resulting in the higher *F*_ST_ between WVX and WVV-YC. Finally, the lowest *F*_ST_ (0.156) was between WVV-YC and WVV-JH, which is still higher than expectation given the short introduction history of WVV-JH. In total, we found the WL was distantly related to both WVV and WVX, WVX was much more closely related to the WVV-JH than to the WVX-YC (Figure 4B).

### *TPS* and *BDH* Genes were Selected based on Population Re-sequencing

To reveal the genetic basis of the difference in the medicinal components of the three AF source (sub)species, we identified 1065 candidate genes in selective sweep regions between WVV and WVX, and 5381 candidate genes for WVV to WL, of which 646 were overlapping (Figure S7 and Table S9-S12). Usually, the pharmacological effect of AF source (sub)species WVV, WVX, WL successively decreases, and terpenes are the main medicinal components of AF^51,52^. GO analysis of 646 candidate selected overlapping genes revealed enrichment of genes involved in terpenoid biosynthesis, such as isoprenoid biosynthetic process, isoprenoid metabolic process, terpene metabolic process, terpene synthase activity (Figure S8), indicating that the selected terpenoid genes were most likely medicinal-related key genes.

In common recognition, the top-geoherbalism of TCM has a great impact on the quality of Chinese medicinal materials. We identified 3,937 selected genes in the comparison of WVV-JH (non-top-geoherb region) *vs*. WVV-YC (top-geoherb region) (Figure S9), and based on gene annotation and studies of homologous genes in *Arabidopsis thaliana* or tomato, we inferred selection on genes involving in terpenoid biosynthesis (e.g., *TPS*: *Wv04G0096, Wv04G0097, Wv06G1862, Wv17G1189, Wv22G0083*) and camphor biosynthesis (e.g., *BDH*: *Wv13G0704, Wv23G0381, Wv23G0382* (Figure 5). Whether these selected *BDH* genes affect gene expression and subsequent camphor abundance in the two populations awaits further experimental validation.

**FIGURE 5.**
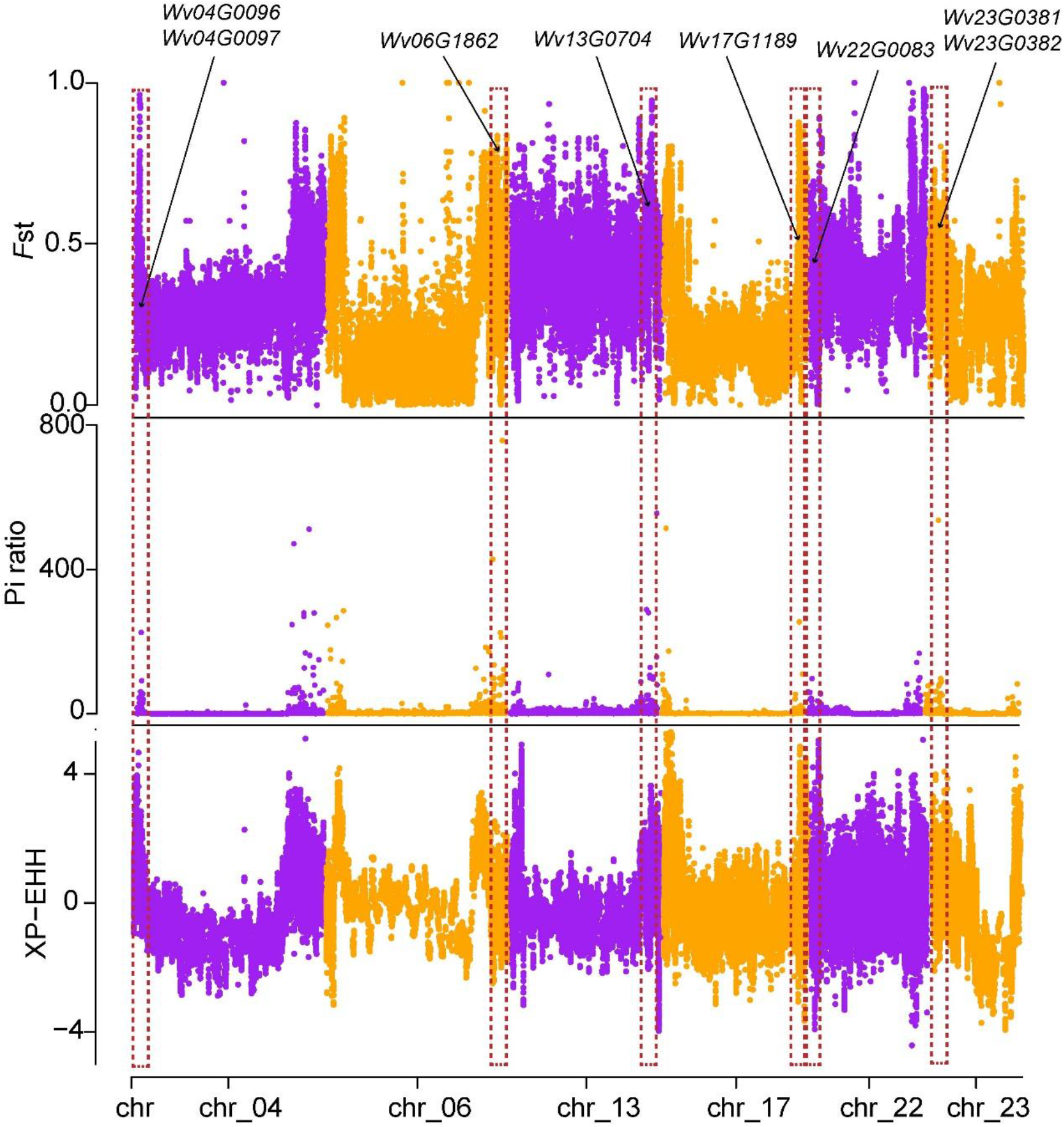
The selective sweep regions identified by at least two statistics among *F*st, π ratio, and XP-EHH methods. Here was comparison of WVV-JH *vs*. WVV-YC. Red dotted boxes and black arrows showed the position of selected genes.

Compared with WVV, WVX has a very low yield and inferior effects, and is often mixed with WVV for sale^53^. To identify molecular markers for the differentiation of the two subspecies, we detected 18,268 nonsynonymous SNPs (involved in 9,565 genes) were fixed in WVV and 707 nonsynonymous SNPs (representing 453 genes) were fixed in WVX (Table S13 and Table S14). These SNPs provided valuable molecular markers to distinguish WVX from WVV.

## DISCUSSION

The high-quality assembly of WVV genome further enriched our understanding of fundamental biology of this species and promoted future comparative genomic, genetic mapping, and gene cloning studies. Our study revealed that the overall expression of related terpenoids biosynthesis genes in WVV was higher than that of WVX, which supplied evidence for the better pharmacological effect of WVV. Meanwhile, we screened ten candidate *BDH* genes that potentially catalyzed borneol into camphor in WVV. *BDH* genes may experience independent evolution after acquiring the ancestral *BDH* genes and followed by subsequent tandem duplications in WVV. Furthermore, from the perspective of whole genome re-sequencing data, four populations including WVV-YC, WVV-JH, WVX and WL are genetically differentiated. The gene flow from WVX to WVV-JH contributed to the increased genetic diversity in the introduced population of WVV in Yunnan (WVV-JH) compared to its top-geoherb region (WVV-YC), which might undergo genetic degradation. Taken together, our study provides new insights into the metabolite biosynthesis, conservation and industrial development of this medicinal material.

Natural borneol, exhibiting better effects than synthetic borneol owing to the higher proportion of (+)-borneol, is a common and valuable composition and widely applied in TCM formulae and daily chemical products for restoring consciousness, removing heat, and relieving pain^10,54-57^. How to quickly and massively obtain natural borneol is a matter of great industrial value. *BAHD* and *BDH* genes take borneol as substrate to produce bornyl acetate and camphor, respectively. Liang et al. has revealed the biosynthetic pathways of bornyl acetate in WVV^27^. However, *BDH*s in WVV have not been studied. We found the tandem duplications of *BDH*s in WVV, *T. chinensis, A. annua*, and *C. camphora*, which possibly increase its expressional dosage and thus elevate the abundance of metabolites^58^. We observed 4 and 14 tandemly duplicated *BDH* genes in WVV on chromosomes 23 and 5, respectively, possibly contributing to the production of camphor. The expressional difference of the selected ten *BDH*s, catalyzing borneol into camphor, and *Wv05G1439, Wv05G1433, Wv05G1432* might account for the difference in camphor abundance in the fruits of WVV and WVX. Further experimental validation is required to confirm their functions.

WVV was originated in Guangdong, and was then transplanted to Yunnan in the 1960s^18^. Interestingly, with such a short cultivation time (∼ 50 years), the Yunnan population was genetically differentiated and demonstrated two-times higher genetic diversity than that of the top-geoherbalism population, which exhibited the lowest genetic diversity. The gene flow from the locally adapted subspecies WVX, which showed the highest genetic diversity among the three (sub)species, contributed to the increased genetic diversity. However, one caveat of our study lies in the absence of comprehensive evaluation of the volatile components in the WVV-YC and WVV-JH when planted in the same environment, which limited our extrapolation about whether the selected *BDH* genes during the introduction is related with its camphor abundance and our inference on the potential pharmacological effects of the two populations. Our results raised the concern for the declined genetic diversity of the top-geoherbalism population with narrow geographic distribution of some medicinal plants. The deteriorated genetic diversity implies species will gradually accumulate deleterious mutations and are more susceptible to serious population shrinkage due to the impact of diseases and insect pests and thus lose the top-geoherbalism advantage^59,60^. At the same time, our study also pointed out the one possible conservation route is to transplant the plants from the top-geoherb region to other suitable habitats with its close wild relatives co-occurring, as the hybridization with the wild relatives with higher genetic diversity will remedy the declined genetic diversity of the species and provide rich genetic resources for the breeding of medicinal plants. Our study calls for the attention to collect and preserve the medicinal plant germplasm resources to enhance the environmental adaptability of TCM^61,62^.

## MATERIALS AND METHODS

### Genome sequencing, assembly and annotation

*Wurfbainia villosa* plants were collected from its top-geoherb regions in Yangchun (111°46′48′′ E, 22°9′36′′ N), Guangdong province, China. Fresh young leaves were selected to extract genomic DNA by using DNeasy Plant Mini Kit (Qiagen, Germany). 50 mg high quality DNA were taken to construct SMRTbell™ libraries and sequenced in circular consensus sequencing (CCS) mode on PacBio Sequell ? platform. Hi-C libraries were constructed from the fresh leaves of WVV, and then sequenced on Illumina NovaSeq 6000 platform^63^.

The genome assembly of WVV by integrating CCS and the Hi-C reads via Hifiasm v0.15.1-r334 with default parameter^64^. Hi-C sequenced reads were mapped to contig level assembly of WVV using Juicer software^65^ and then 3D-DNA pipeline^64^ were used to correct mis-joins, orientation and order, and generate a draft chromosome assembly. Finally, the draft assembly was visualized in Juicebox Assembly Tools (https://github.com/aidenlab/Juicebox) and conducted manual correction to obtain chromosome-level genome of WVV. Its completeness was evaluated by BUSCO v5.1.2^66^.

EDTA pipeline^67^ was employed to identify TE in the WVV genome, and MAKER2 pipeline^68^ was applied to predicate coding gene structure from *ab initio* predictions, homolog proteins and transcriptome data. Functional annotations of coding sequences were aligned by BLASTP (“-e-value 1e–5”) in Swiss-Prot databases and annotated using online EGGNOG-MAPPER (http://eggnog-mapper.embl.de/) for Pfam, GO and KEGG.

### RNA Sequencing

Plant materials of WVV, including rhizomes (Rh), young leaves (LY), flowers (F), fruits of 10-day after flowering (P1), 30-day after flowering (P2), 60-day after flowering (P3), 90-day after flowering (P4), and WVX, including rhizomes (XRh), young leaves (XLY), fruits of 75-day after flowering (XP1), 90-day after flowering (XP2) were used for RNA sequencing and three biological replicates for each sample.

Total RNA was extracted using RNAprep Pure Plant kit (TIANGEN, China) and 20 mg RNA was used for reverse transcription to synthesize cDNA. RNA sequencing was performed on Illumina NovaSeq 6000 platform. All the clean reads were mapped to WVV genome using HISAT2 software and the transcripts per million reads (TPM) was calculated using counts from featureCounts for finally used to measure the expression level^69,70^.

### Transcriptional Regulation of Bornyl Acetate and Camphor Biosynthesis

To identify transcriptional regulatory networks between bornyl acetate, camphor biosynthetic genes and TFs, a series of gene expression and co-expression network analysis were performed. Differentially expressed genes (DEGs) in different tissues were employed to construct a co-expression network using weighted gene co-expression network analysis (WGCNA)^71^. The co-expression network modules were attained and PlantTFDB were used with default parameters to identify TFs in the WVV genome^72^. The networks between genes and TFs were visualized in Cytoscape^73^.

### Phylogenetic Analysis of *BDH* Genes Which Catalyzed Borneol into Camphor

To identify candidate *BDH* genes which could convert borneol into camphor, a total of 116 homologous proteins were identified through querying in WVV, *C. camphora, S. officinalis, T. chinensis, A. annua*, and *R. officinalis*. Then combined with nine experimentally validated *BDH* enzymes previously, the *BDH* phylogenetic tree was constructed and iTOL was used to visualize and edit the tree^74^.

### Re-sequencing and Variant Calling

DNAs from 39 leaf tissue, which contained 20 samples of WVV (10 from Yangchun, Guangdong and 10 from Jinghong, Yunnan), 10 WVX and nine WL, were used to construct the 150-bp paired-end libraries in BENAGEN (Wuhan, China), and then were sequenced with the DNBSEQ-T7 platform (MGI, China).

The re-sequencing data of 39 accessions initial quality control was performed by FastQC^75^ and adapters were removed by Trimmomatic^76^. The treated data were then mapped to WVV genome using BWA-MEM^77^. The mapped reads were sorted using Samtools to generate bam files^78^. After that, duplicates were removed using Picard, and individual gvcf files were produced by using GATK (v4.0.12) HaplotypeCaller^79,80^. Finally, we used GATK CombineGVCFs to the gvcf files were combined using to obtain raw vcf files and the SNPs were hard filtered using GATK VariantFiltration (QD<2.0 ∥ QUAL < 30.0 ∥ MQ < 40.0 ∥ FS > 60.0 ∥ SOR > 3.0 ∥ MQRankSum < -12.5 ∥ ReadPosRankSum < -8.0), and only biallelic SNPs were selected for further analysis.

### Population Genetic Diversity and Structure Analysis

To infer the basal group of AF origin plants, we constructed a phylogenetic tree based on 160,967 filtering SNPs (MAF ≥ 0.05, missing rate ≤ 0.1 and minimum distance between two SNPs ≥ 1 kb). We calculated the p-distance matrix of the 39 accessions with VCF2Dis (https://github.com/BGI-shenzhen/VCF2Dis), and the matrix was used to build the neighbor-joining (NJ) tree. PCA was performed with plink^81^ and the population structure was analyzed using the STRUCTURE software^82^ with the likelihood of ancestral kinships (*K*) from 3 and 4, both of used SNPs were filtered.

Nucleotide diversity (θ_*π*_) was determined for WVV-YC, WVV-JH, WVX, and WL population using VCFtools^83^ with parameters: 100-kb sliding window and 50-kb step size. We calculated genetic differentiation (*F*_ST_) among different groups using the same method.

### Detection of Sweeps

To avoid bias due to potential gene flow between WVV-LH and WVX, we conducted selective sweep analysis excluding admixed samples as identified by the structure analysis. We calculated the XPEHH^84^, *F*_ST_ and π value of each SNP, *F*_ST_ and π value based on a sliding window of 10-kb and a step size of 1-kb. Regions ranked top 5% of the score in any two of the methods were defined as putative selective sweeps.

## DATA AVAILABILITY STATEMENT

The data presented in the study were deposited in National Center for Biotechnology Information (NCBI), and accession number was PRJNA910288.

## CONTRIBUTIONS

LW, DY, XC, SS, and XH conceived and designed the study. XC, DY, LW, LZ and JY prepared the materials. JJ performed flow cytometry experiment. XC, SS, XH, CL, BN, and ZH performed data analysis. XC, SS, XH and LW wrote the manuscript. LW, SS, XH and DY revised the manuscript. All authors read and approved the final draft.

## ACKNOWLEDGMENTS

This study was supported by Yunnan Science and Technology Talents and Platform Program (Academician and Expert Workstations, 202205AF150071), the National Key Research and Development Program of China (Nos. 2020YFA0907900, 2022YFD1600300, and 2017YFC1701100), Open Projects of Guangxi Key Laboratory of Medicinal Resources Conservation and Genetic Improvement (No. KL2022KF01), the Shenzhen Science and Technology Program (No. KQTD2016113010482651), special funds for Science Technology Innovation and Industrial Development of Shenzhen Dapeng New District (Nos. RC201901-05 and PT201901-19), the China Postdoctoral Science Foundation (No. 2020M672904), the Basic and Applied Basic Research Fund of Guangdong (No. 2020A1515110912), Scientific and Technological Talents and Platform Plan (Academician and Expert Workstations, 202205AF150071) and National Natural Science Foundation of China (Nos. 32070242 and 82260736).

## SUPPLEMENTARY INFORMATION

The Supplementary Material for this article can be found online:

TABLE S1 | Pseudochromosome and scaffold length of the genome.

TABLE S2 | Statistics of genes annotation in WVV.

TABLE S3 | The statistical results of gene functional annotation.

TABLE S4 | TE sequences of the genome.

TABLE S5 | ID of BDHs used in this study.

TABLE S6 | Gene expression levels (TPM) in different tissues and fruit developmental stages in WVV and WVX.

TABLE S7 | Interaction genes corresponding to TFs of nine *BAHD* genes in WVV.

TABLE S8 | Interaction genes corresponding to TFs of ten *BDH* genes in WVV.

TABLE S9 | The selected genes in the comparison of WVV-JH *vs*. WVV-YC.

TABLE S10 | The selected genes in the comparison of WVV-YC *vs*. WVV-JH.

TABLE S11 | The selected genes in the comparison of WVV *vs*. WVX.

TABLE S12 | The selected genes in the comparison of WVV *vs*. WL.

TABLE S13 | The nonsynonymous SNPs which fixed in WVV genome.

TABLE S14 | The nonsynonymous SNPs which fixed in WVX genome.

FIGURE S1 | Hi-C interaction maps for WVV genome.

FIGURE S2 | The ML tree of *TPS* genes in WVV and the functionally verified proteins in the previous study.

FIGURE S3 | TFs potentially regulating nine *BAHD* genes.

FIGURE S4 | TFs potentially regulating ten candidate *BDH* genes.

FIGURE S5 | Principal component analysis of four populations.

FIGURE S6 | The optimal number *K* of populations in the STRUCTURE analysis.

FIGURE S7 | The Venn diagram of the selective sweep regions.

FIGURE S8 | GO analysis of 646 overlapped genes under selection.

FIGURE S9 | GO analysis of selected genes between WVV-YC and WVV-JH.

Supplementary data 1: Amino acid sequences of BDH used in this study.

## Conflict of Interest

The authors declare that the research was conducted in the absence of any commercial or financial relationships that could be construed as a potential conflict of interest.

---

**FIGURE S1.**
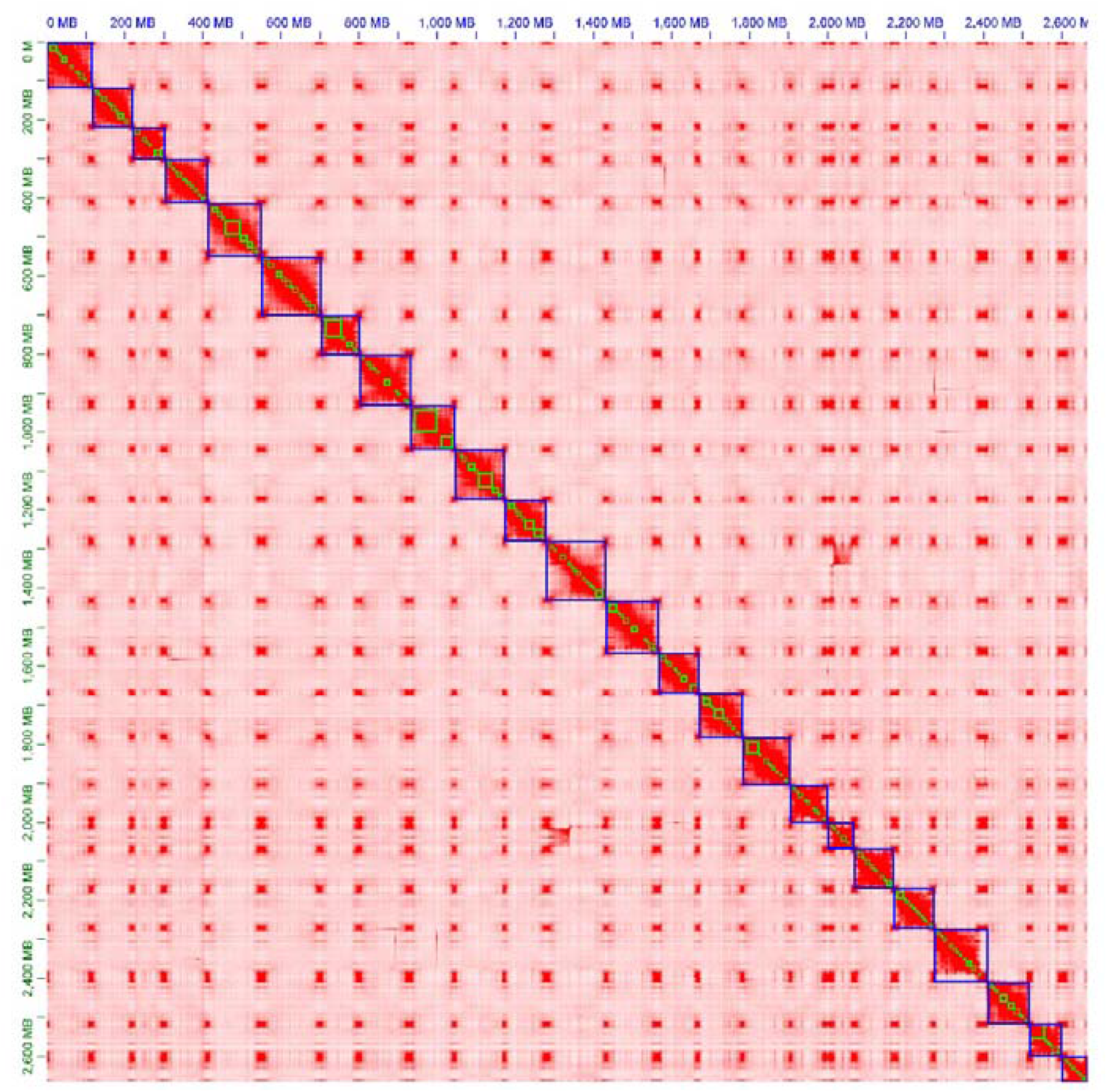
Hi-C interaction maps for WVV genome.

**FIGURE S2.**
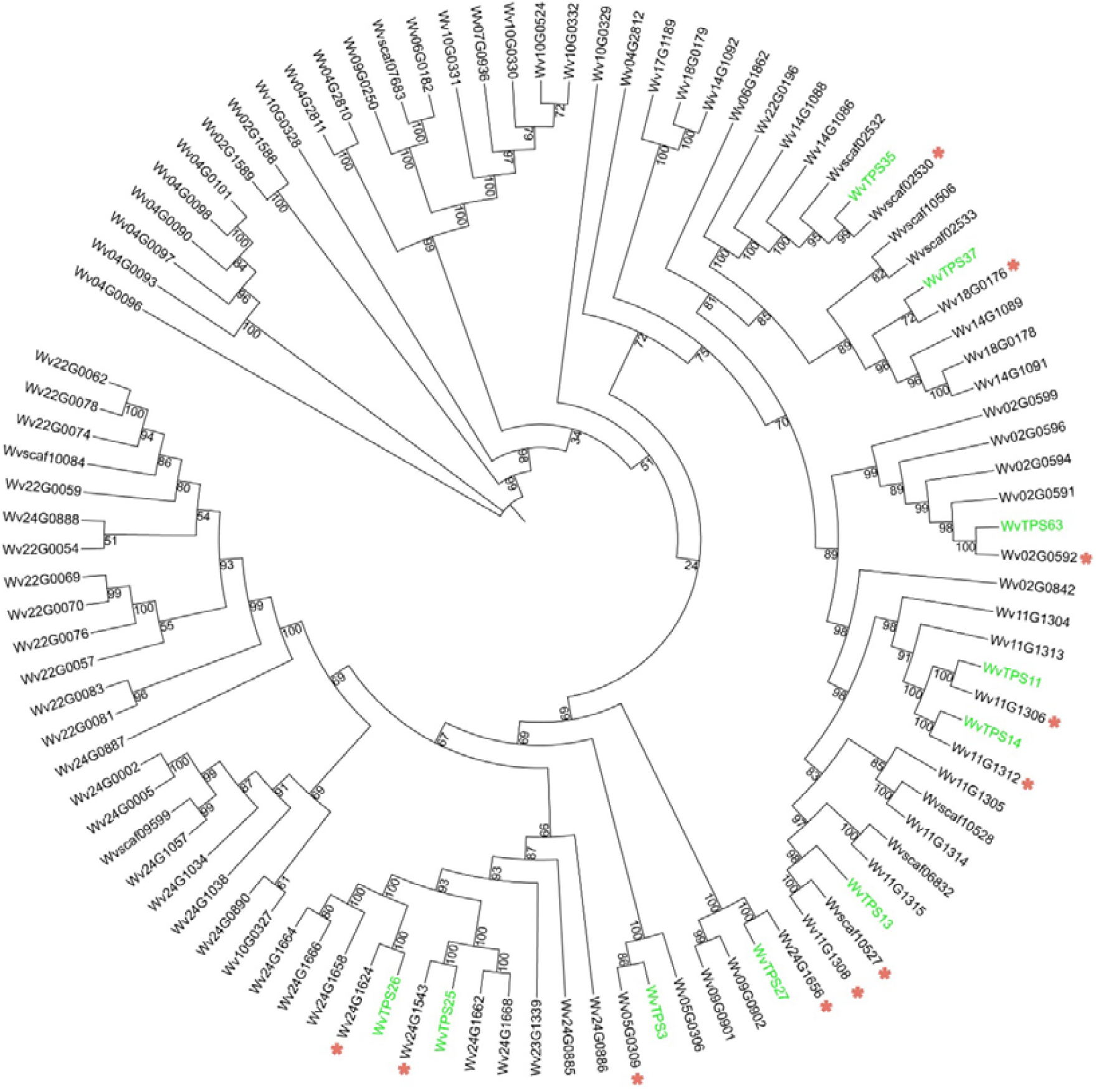
The ML tree of *TPS* proteins in WVV and the functionally verified proteins in the previous study^17^. The genes in green font indicated the ones verified in previous researches, and the red asterisks showed the corresponding genes with the smallest phylogenetic distance.

**FIGURE S3.**
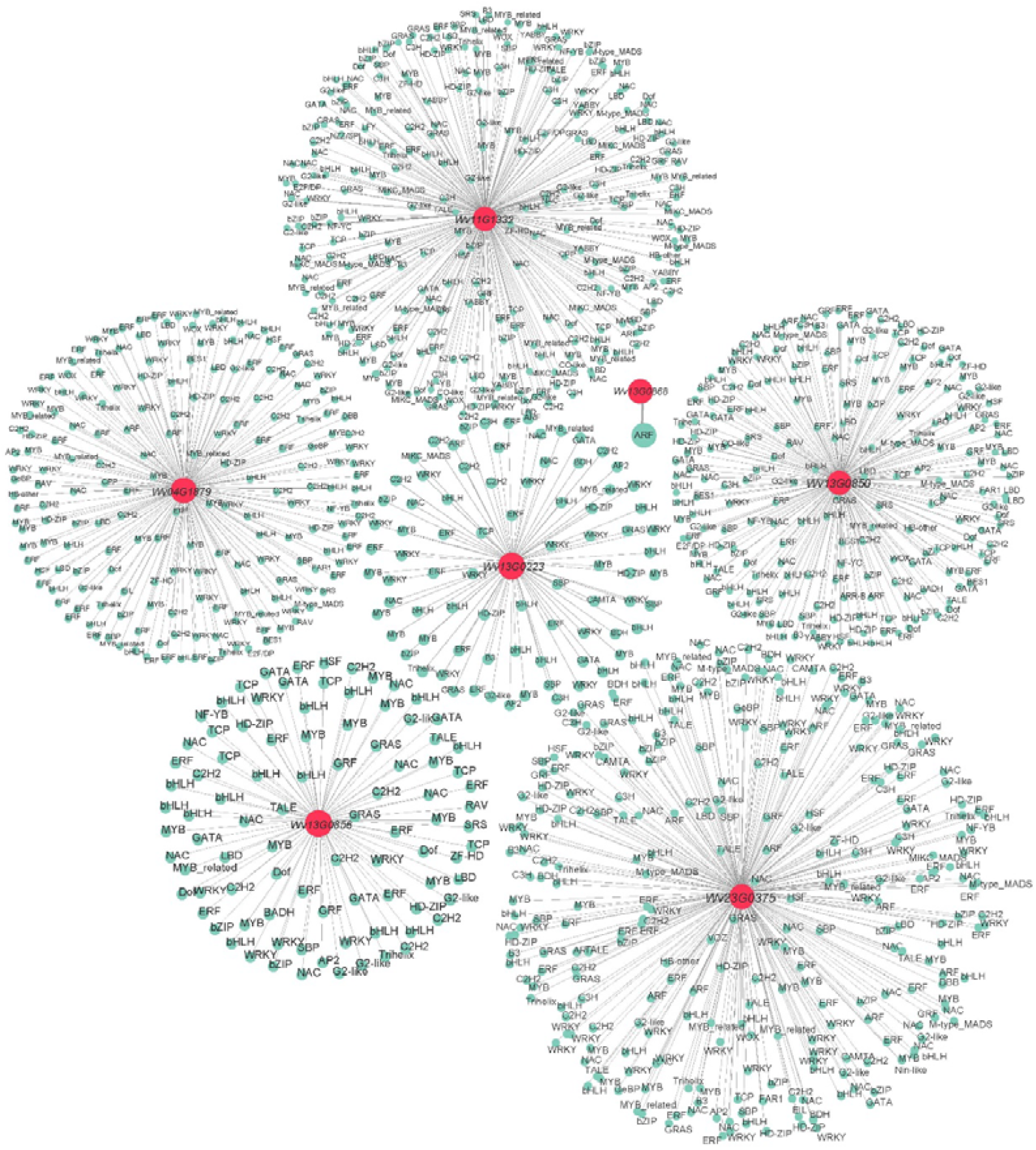
TFs potentially regulating nine *BAHD* genes. The red dots are nine *BAHD* genes, and the green dots connected to them indicate the TFs that may be involved in the regulation of *BAHD* genes.

**FIGURE S4.**
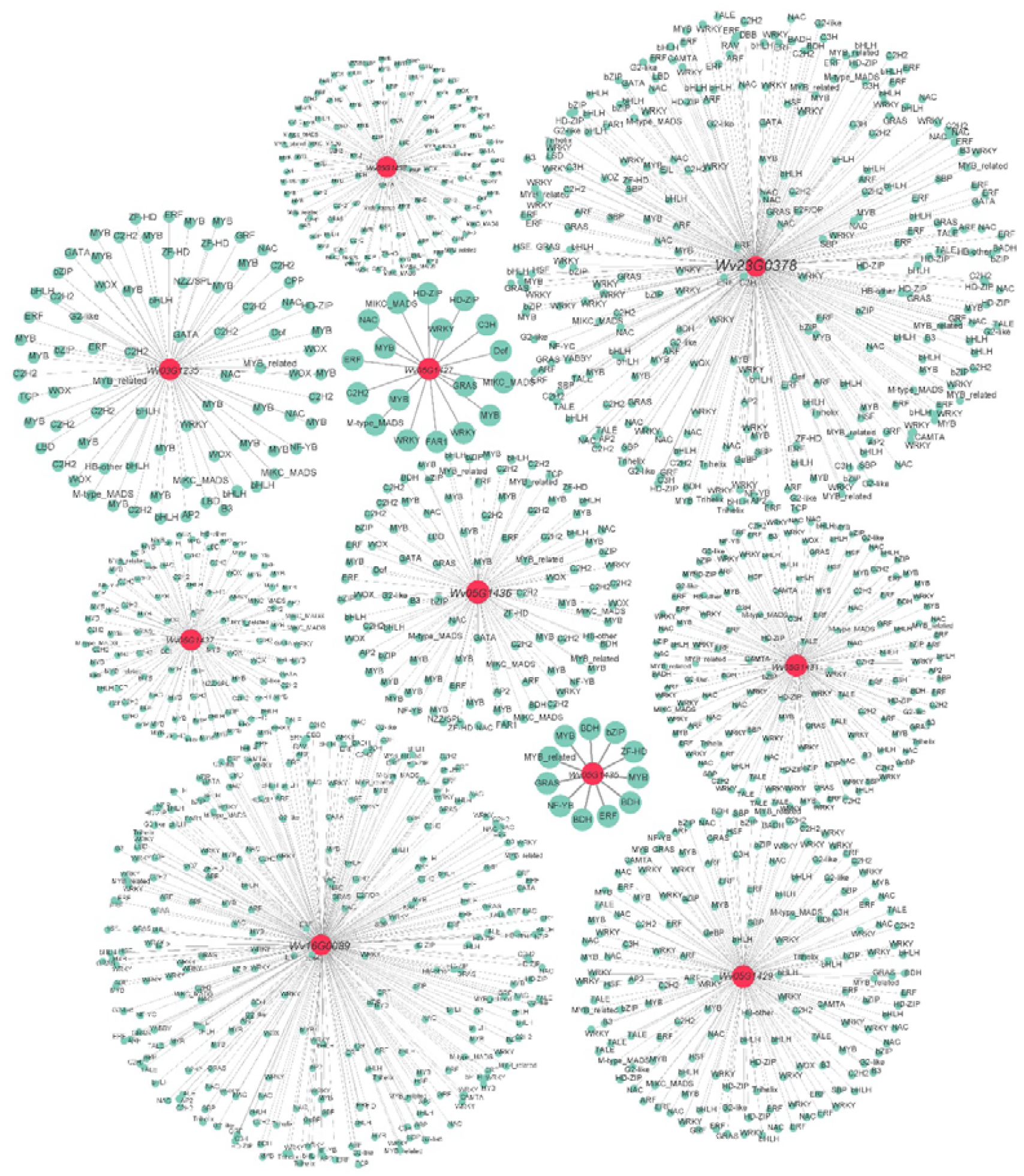
TFs potentially regulating ten candidate *BDH* genes. The red dots are ten *BDH* genes, and the green dots connected to them indicate the TFs that may be involved in the regulation of *BDH* genes.

**FIGURE S5.**
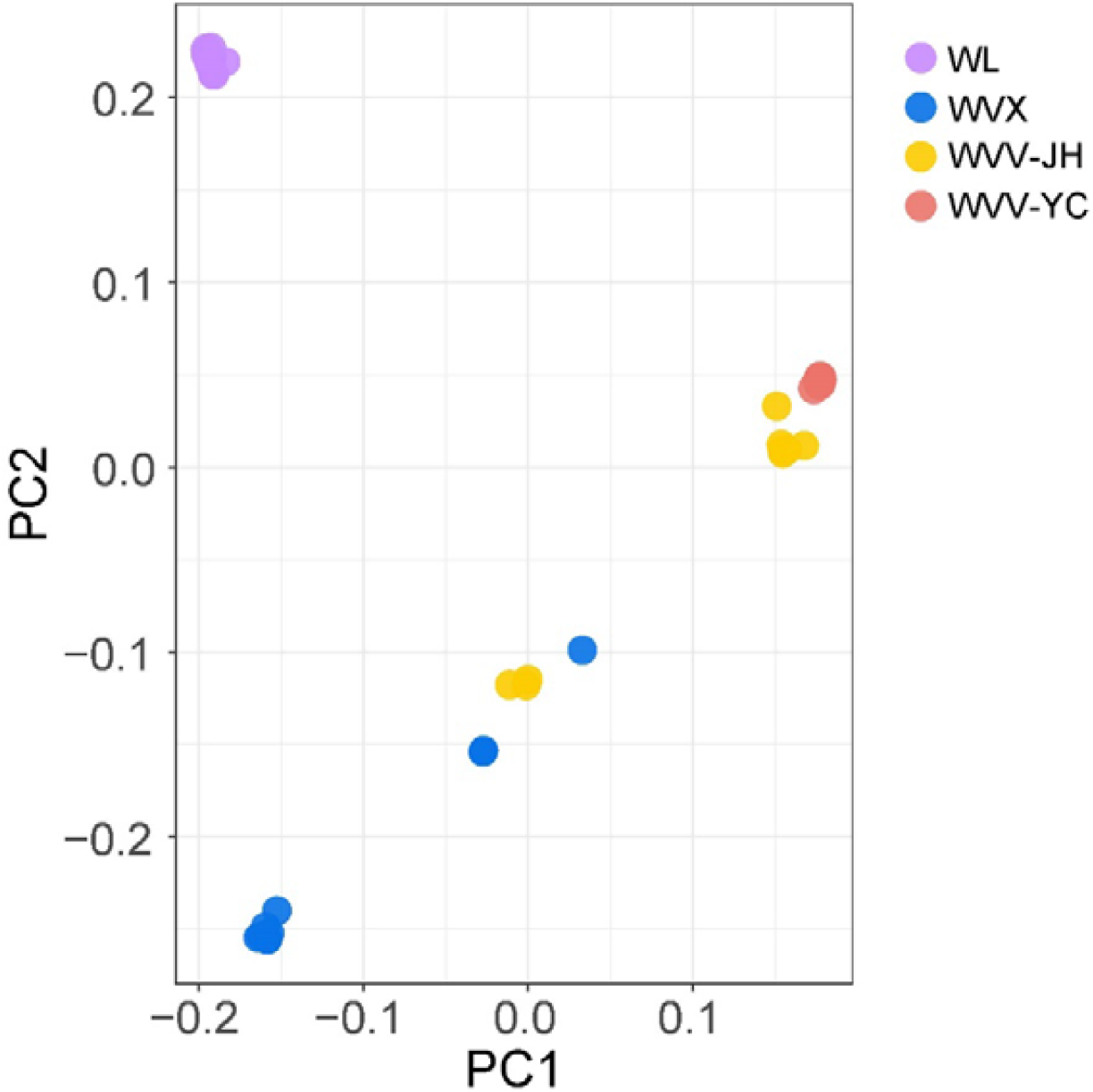
Principal component analysis of the four populations.

**FIGURE S6.**
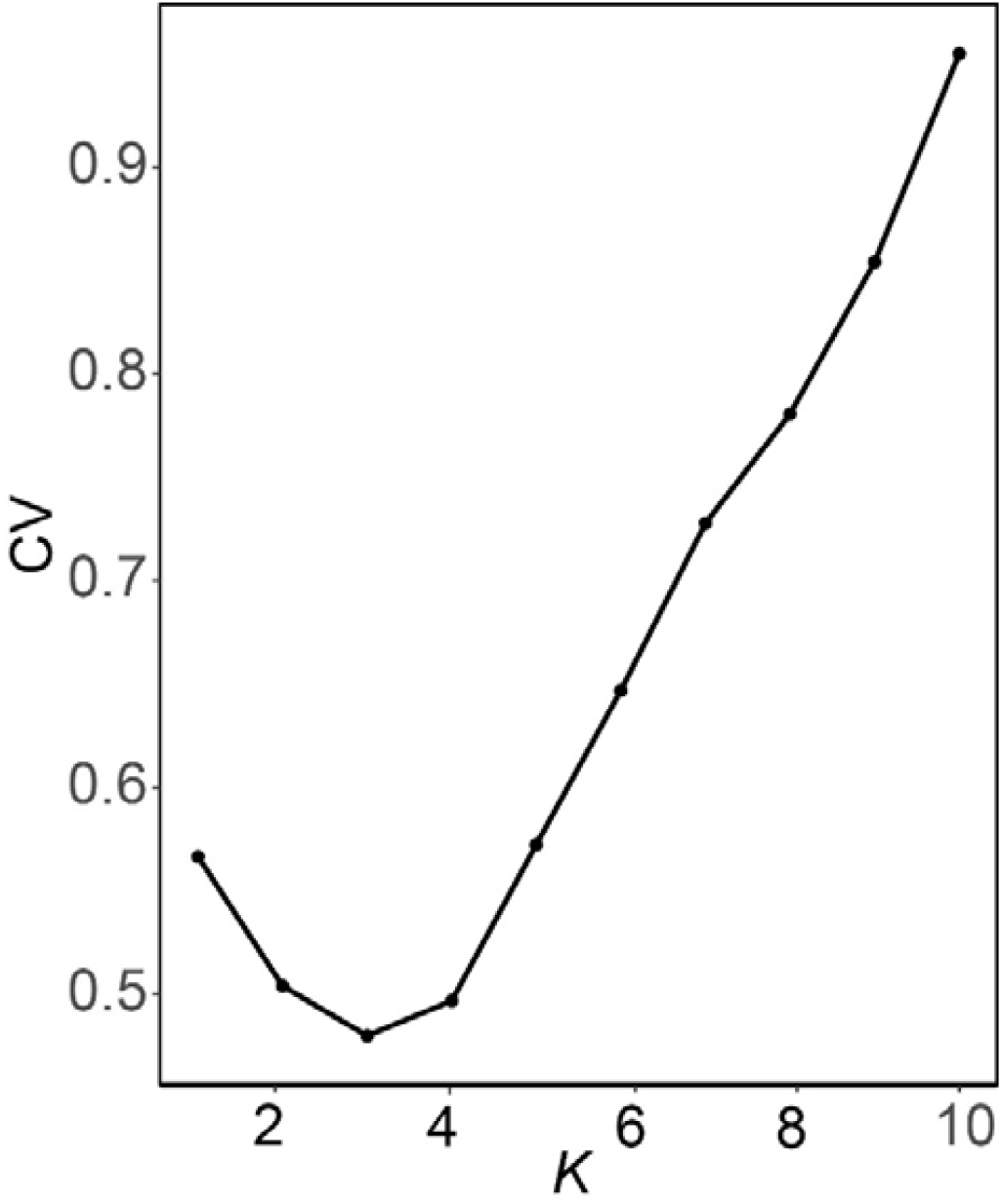
The optimal number *K* of populations in the STRUCTURE analysis. The y-axis CV showed Cross-Validation.

**FIGURE S7.**
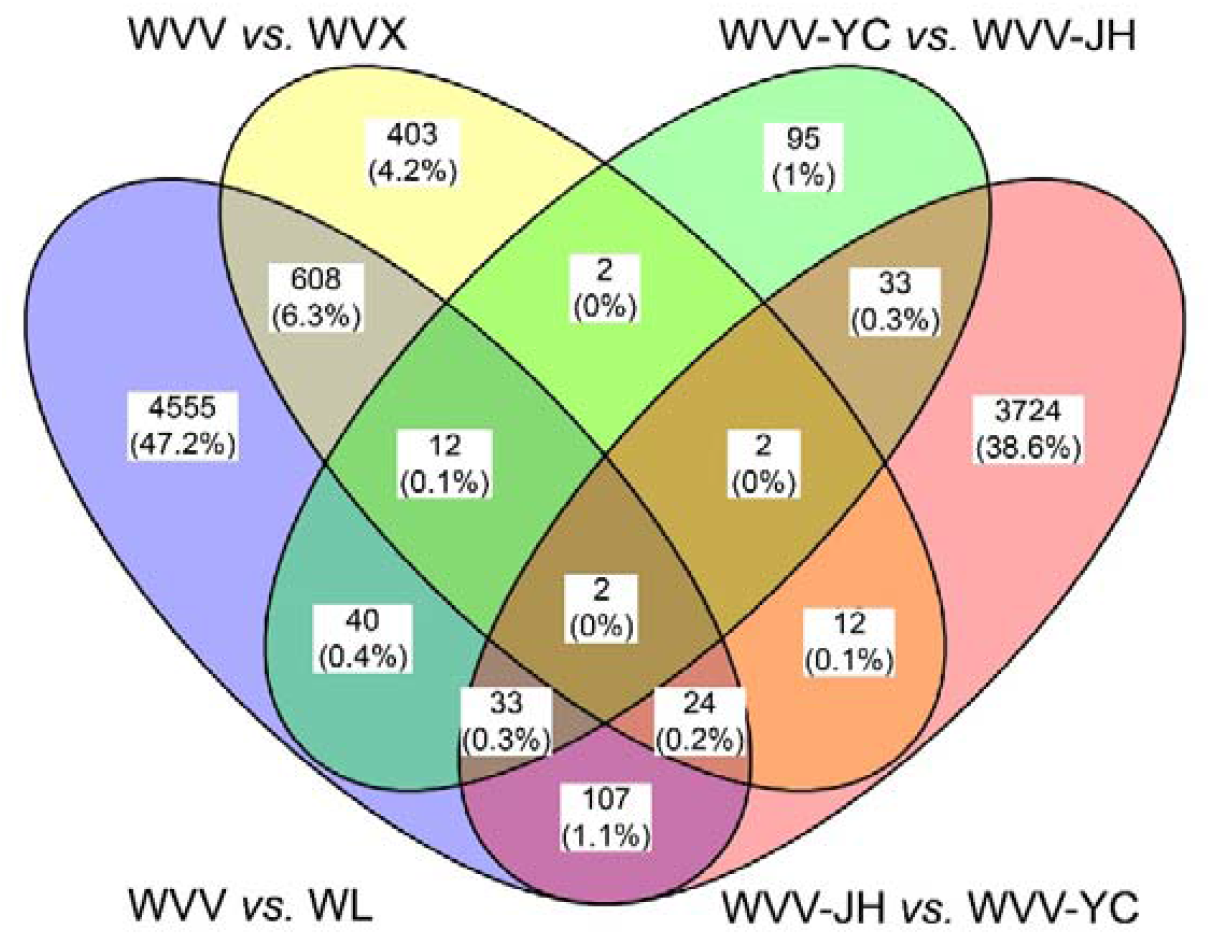
The Venn diagram of the selective sweep regions. Numbers on different color blocks showed the numbers of candidate genes in selective sweep regions. Percentages represented proportions in the union of the four comparisons.

**FIGURE S8.**
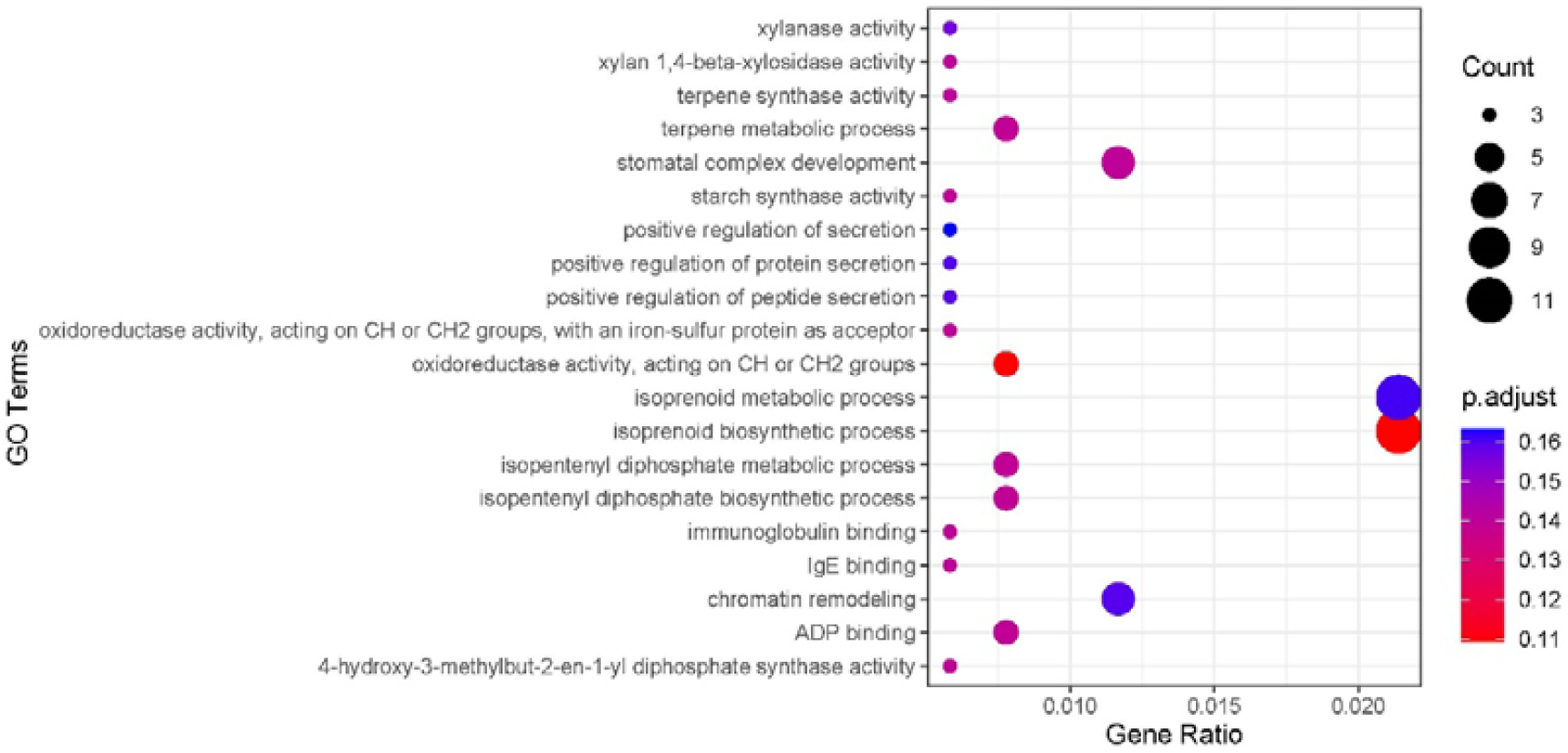
GO analysis of 646 overlapped genes under selection. The y-axis represented GO terms.

**FIGURE S9.**
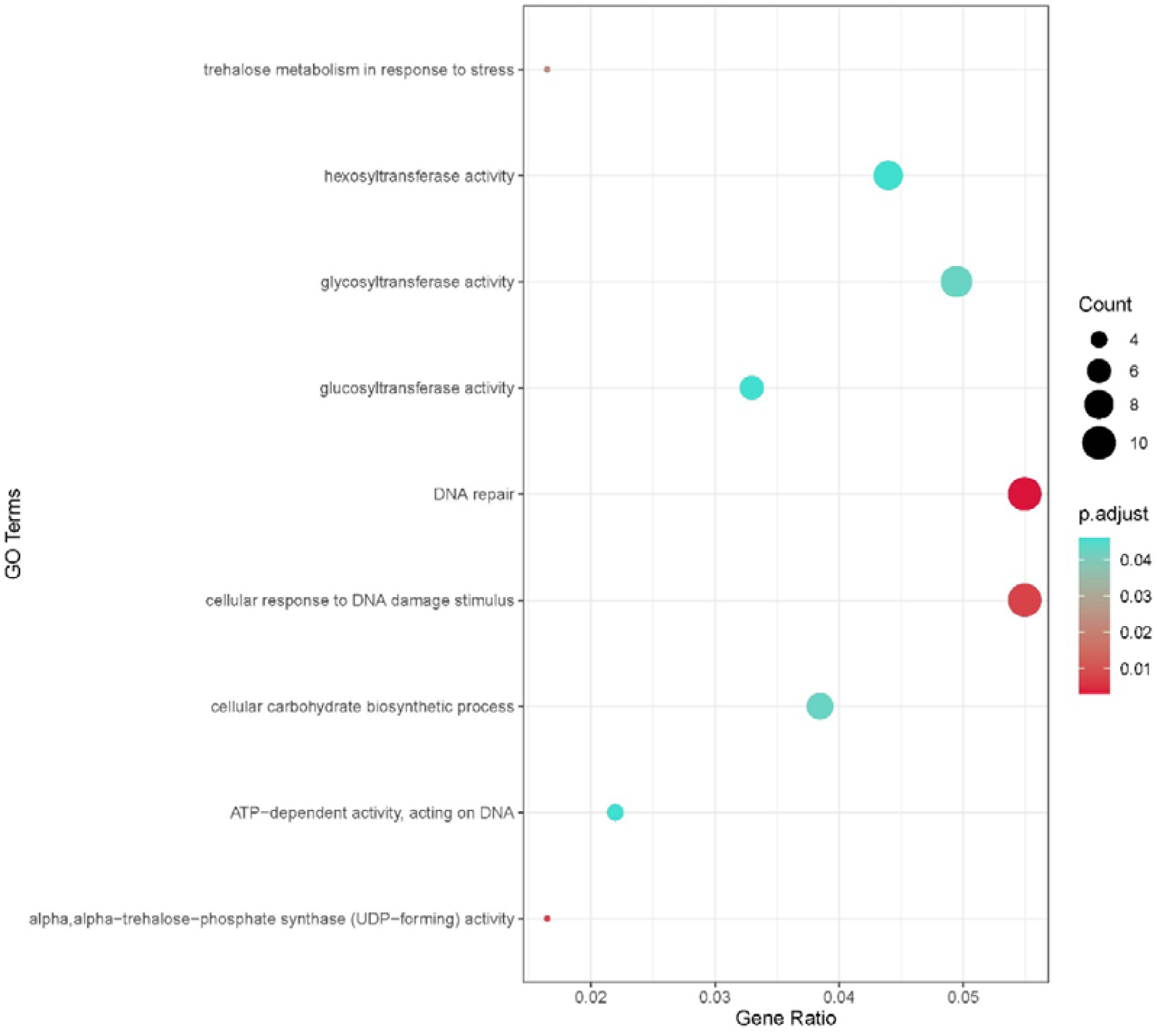
GO analysis of selected genes between WVV-YC and WVV-JH. The y-axis showed GO terms.

## Notes

### Competing Interest Statement

The authors have declared no competing interest.

